# Spinal Motor Neuron Pools May be Partly Driven by Impulsive Common Inputs

**DOI:** 10.1101/2025.08.13.670065

**Authors:** Javier Yanguas Mayo, Alejandro Pascual Valdunciel, Stuart N Baker, Pablo Laguna, Dario Farina, Jaime Ibáñez Pereda

## Abstract

Spinal motor neurons serve as the link between the nervous system and muscles. As the final common pathway of the neuromuscular system, they receive inputs from both higher-level controllers and afferent pathways. It is often assumed that spinal motor neurons are primarily driven by continuous common inputs (cCI) within different frequency bands. Within this framework, the motor neuron pool behaves as a linear amplifier of the cCI. However, this framework overlooks the possibility that motor neurons could also be driven by impulsive common inputs (iCI), which can induce synchronization among them and disrupt the linear transmission of other synaptic inputs at the pool level. To test this hypothesis, computational simulations and experimental data from human subjects were used to characterize different aspects related to motor neuron spiking synchronization at the pool level. Our findings suggest that, indeed, iCI can account for relevant features observed in experimental data such as the presence of synchronization events at the pool level. We also observed that such impulsive inputs can affect the linearity in the transmission of cCI by the motor neuron pool. This study represents pioneering indirect evidence of the existence of iCI as inputs to motor neurons.

**Significant Statement:** Motor unit pool behavior in terms of spiking synchronization and spectral content typically observed in experimental recordings cannot be reproduced in simulations that only use continuous common inputs (cCI) to motor neurons. This study shows, for the first time, evidence supporting that spinal motor neurons receive a portion of their synaptic input in the form of impulsive common inputs (iCI) that synchronize their activity. The study also shows how such iCI can affect the linear transmission of other cCI by the motor neuron pool.

## Introduction

Spinal Motor neurons (MNs) integrate inputs received from different parts of the nervous system and transmit this information to the muscles Gandevia (2001). MNs can be modeled as elements receiving two types of inputs: those shared across a pool of MNs (common inputs or CI) and those that are different for each MN (independent inputs or II) De Luca and Erim (1994); Farina et al. (2014). During constant-force muscle contractions, spinal MNs fire at steady rates. In this context, the power spectral density of the MN activity is largely dominated by the average discharge rate of the MNs and its harmonics. The power related to the inputs received by the MN (especially at frequencies above the discharge rate) are less prominent Nakao et al. (1997). This observation is explained by the behavior of individual MNs as non-linear systems. However, when the activity of groups of MNs receiving common inputs is summed, the CI are amplified proportionally to the total number of MN discharges. This characteristics of the MN pool to act as a linear amplification system of CI has been previously described and it applies to both low-and high-frequency inputs received by the MNs Baldissera et al. (1998); Farina et al. (2014); Farina and Negro (2015). The implication of this framework is that the summed activity of a sufficiently large number of MNs may be considered a reliable estimate of the net CI they receive. Since there exists evidence of stable transmission of brain signals to the MNs, this also implies that the compound MN activity may be used to estimate CI from the brain Ibáñez et al. (2021).

The conceptual framework of MN pool acting as a linear amplifier of CI is valid only for continuous CI (cCI). This may in part be due to the intuition that, for relatively steady forces, the neural inputs to MNs will also have slow continuous dynamics. However, there are many examples in the literature suggesting that a portion of the CI that MNs receive could be better modeled as a series of impulses or short-lived bursts. For example, intermittent submovements have been observed during slow-varying contractions Karniel (2013); Susilaradeya et al. (2019), and brief inhibitory-excitatory inputs to MNs have been detected when passively attending to external stimuli Novembre et al. (2018). The frequently observed phenomenon of cortical beta bursts transmitted to the MN pool Echeverria-Altuna et al. (2022) is a further example of intermittent inputs. These types of inputs may be sufficiently strong to align the firing times of different MNs transiently. This transient synchronization will impact on the linear behavior of the MN pool: increased spike alignment across MNs will make the summed activity of the pool more similar to that of a single MN in terms of spectral distribution of power (with a dominance of the spectral peaks at the discharge rate and harmonics).

The existence of impulsive CI (iCI) projecting to MN pools and their impact on MN behavior has not been systematically addressed. As these CI are brief and intermittent, their effects on MN activity are also restricted in time (short-lived events of synchronization). Importantly, standard methods used to analyze muscle signals are either based on the analysis of pairs of motor units or assume stationarity of the analyzed signals. These methods are therefore not suitable to study non-stationary impulsive signals, which may explain why the existence of such iCI has not been analyzed so far.

Here, we analyze experimental data from humans and compare it with computational simulations to verify whether MN pools receive iCI and if these inputs alter the linearity assumption in CI transmission by MN pools. Experimental results indicated that MN pool activity presents events of high synchronization at the population level, which agrees with simulation results testing the hypothesis that MN pools receive iCI. Additional tests also showed that the iCI alter the linear behavior of the MN pool, affecting the linear transmission of cCI and therefore their estimation based on the measurement of motor unit activity.

## Materials and Methods

The primary aim of this study was to determine whether the activity of the MN pool is partly driven by impulsive-like input components. For this purpose, we combine the analysis of both experimental data from human subjects and data of three simulated scenarios that aimed to reproduce the experimental data. The next subsections describe the characteristics of the experimental data analyzed, the computational model and simulations used, and the procedures used to derive the results of the study.

### Human data

The characteristics of the experimental human data analyzed in this study have been described in a previous study Ibáñez et al. (2021). All subjects provided written informed consent. The study was approved by the University College London Ethics Committee (Ethics Application 10037/001) and was carried out in accordance with the Declaration of Helsinki. The experimental procedures are briefly summarized below.

The experiments involved the recording of the tibialis anterior (TA) muscle EMG electrical activity of 19 subjects (two females), during two consecutive repetitive experiments. In this study we only considered a subset of the subjects based on the number of motor units that could be decomposed from the surface EMG recordings. This is due to the fact that we needed sufficiently large populations of motor units decomposed so that the metrics based on motor unit synchronization (described in the next sections) were reliably estimated. Therefore, we included only subjects with at least 15 MUs decomposed from the muscle, resulting in a total of 14 subjects. Also, due to the difficulty to track the same MUs across different recording experiments, we only used one of the two recorded experiment per subject. For consistency, we selected the experiment from which we could identify more MUs in each subject.

The recorded EMG was obtained while subjects sat on a straight-backed chair with their knees flexed at 90° and their right foot positioned beneath a custom-made lever designed to measure ankle dorsiflexion forces during isometric contractions. Ankle force measurements, along with high-density electromyography (HD-EMG) recordings of the TA (using grids of 5×13 channels with 8-mm inter-electrode distance), were recorded using a multichannel EMG amplifier (Quattrocento, OT Bioelettronica). A sampling rate of 2048 Hz was used.

After a standardized warm-up, subjects performed three maximal voluntary isometric contractions (MVCs) of the recorded muscle in each session, with at least 30 seconds of rest between trials. During MVCs, they were instructed to contract as forcefully as possible for at least 3 seconds. The highest recorded force across the three trials was taken as the final MVC value and digitally stored. Subsequently, subjects followed ramp-and-hold force trajectories displayed on a monitor. Each trial consisted of a 2-second ramp contraction from 0% to 10% MVC, followed by a 60-second holding phase at 10% MVC. During this phase, subjects maintained an isometric force at 10% MVC. Each subject repeated the ramp-and-hold contraction twice, with a 2-minute interval between trials.

### Simulated Data

In this section, we present the computational model used to simulate the activity of a MN pool and describe each simulation conducted with the model.

### Computational Model Overview

The model used for the simulations is the one presented in Williams and Baker (2009). In short, the model simulates a MN pool consisting of 177 slow-type MNs. This represents the type of MN active during low-level muscle contractions. Each of the *j*-th MNs simulated, *j ∈* {1, …, 177} is represented as a conductance-based two-compartment model (dendritic and somatic compartment), following Hodgkin-Huxley kinetics. It includes 8 active conductances found in mammalian MN (somatic: g_Na_, g_K-dr_, g_Ca-N_, g_K(Ca)_ and g_NaP_, dendritic: g_Ca-L_, g_Ca-N_ and g_K(Ca)_). The membrane potential change in the somatic compartment is determined by:

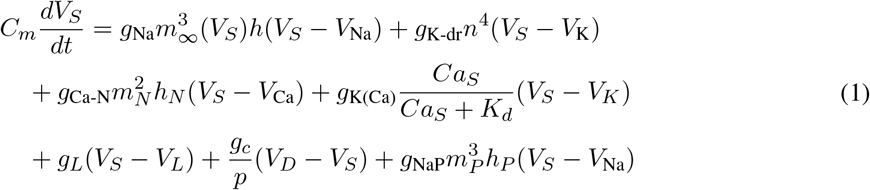

The membrane potential change in the dendritic compartment is determined by:

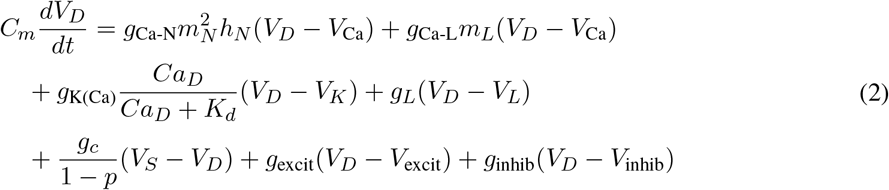

where *V*_*s*_ and *V*_*d*_ are the voltage in the soma and dendrite compartments,*p* is the ratio of somatic surface area to total cell surface area, which was held constant in this study. The gating variable *m* is the sodium-channel activation gate, the gating variable *h* is the sodium channel inactivation gate and the gating variable *n* is the potassium channel activation gate. Each gating variable is governed by an equation of the form:

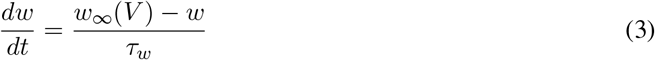

Where *τ*_*w*_ is the activation and inactivation time constant, and *w*_*∞*_(*V*) is the steady-state activation and inactivation function for the gating variable *w*, and it is given by:

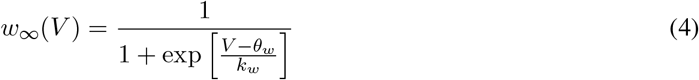

where *θ*_*w*_ is the half-voltage activation and inactivation function for the gating variable *w*. Similarly, *k*_*w*_ is the activation and inactivation sensitivity for the gating variable *w*. The parameters used to solve the above equations were the same as the ones used in Williams and Baker (2009).

The model included a population of 64 Renshaw Cells (RC) that constitute the only inhibitory input to the MNs. Each MN received projections from 20 RC and each RC received projections from 50 MN. The membrane potential of the RCs also followed Hodgkin-Huxley kinetics. The RC had a mean firing rate of 11 spikes/s. The computational model also simulated force, as described in Williams and Baker (2009). In this model, the motor unit force was scaled by a gain factor, which depended on the unit’s twitch contraction time and the interspike interval. Details can be found in the original article.

The sampling rate of the computational model was 5000 Hz. The differential equations governing MN membrane potential were solved using the exponential integration scheme MacGregor (2012), with a time step of 0.2 ms. The model was coded and run in MATLAB (version 2024a, The Mathworks Inc., USA).

#### Simulation scenarios

In all the simulations, each of the 177 MNs was modeled as receiving a CI, *m*_CI_(*t*), composed of several components, and an independent input, *m*_II_(*t*), that was different for each MN. Therefore, the total input to each MN was defined as *m*(*t*) = *m*_II_(*t*) + *m*_CI_(*t*).

The CI components depended on the simulated scenario, as explained below. The independent inputs *m*_II_(*t*) were simulated as a constant value 𝒞_*j*_ plus a MN-specific individual runs of white Gaussian noise, with zero mean and a standard deviation equal to the MN-specific 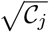 value, quarantining a spike variability within the mean interspike range. In this study, *𝒞*_*j*_ is a constant value whose statistics across MNs determined the MN average discharge rate (DR). We fitted the model to capture the broad variability in MN firing observed in our experimental dataset. To do so, the firing rate of each MN was individually adjusted by varying the value of *𝒞*_*j*_, the constant current. In the simulated MN pool, the slowest MN exhibited a mean DR of 7.3 Hz, while the fastest reached a mean DR of 13.4 Hz. The overall mean DR across the MN pool was 10.4 Hz, with a standard deviation of 1.4 Hz. This average aligns with previously reported values for low-level contractions of the TA muscle Ibáñez et al. (2021). For each simulation scenario, we ran 10 simulations of 30 s each, unless otherwise stated. The results were averaged across the simulations. The different simulation scenarios used in this study are summarized in Table 1 and described below.

**Table 1:** Simulation scenarios as a function of the types of CI to the MN pool included in the simulation.

| Scenario MN Inputs | Noisy<br>II | Constant<br>II | cCI | iCI |
| --- | --- | --- | --- | --- |
| iCI | x | x | - | x |
| cCI | x | x | x | - |
| Control | x | x | - | - |
| Combined CI | x | x | x | x |

Figure 1 represents graphically the simulated scenarios and the simulated inputs to the MN pool.

**Figure 1.**
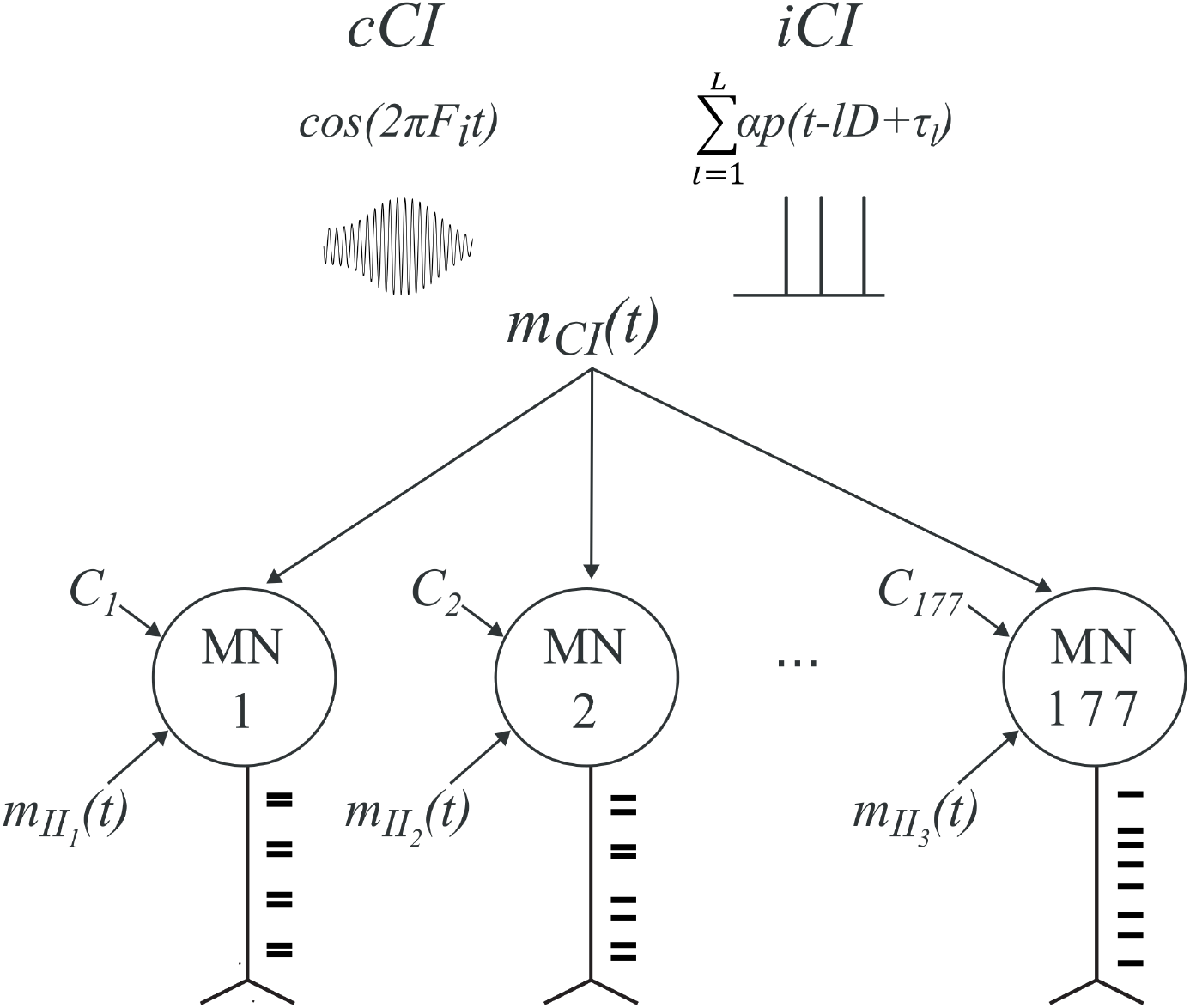
Graphical representation of the computational model and the simulated scenarios. 𝒞 _j_ represents a constant input, different for each j-th MN. m_CI_(t) represents a common input for the MN pool that modulates the activity of the MNs. In our work, m_CI_(t) was simulated as a sinusoid at a particular frequency (cCI) or as a series of impulsive pseudo-periodic bursts (iCI). m_II_(t) represents the independent input, different for each MN.

#### iCI scenario

The main hypothesis of this study is that the MN pool receives iCI components that synchronize the activity of individual MNs with each impulsive event.

To analyze this hypothesis, we defined this CI component as a train of excitatory rectangular bursts (except during the initial and final one-second intervals), 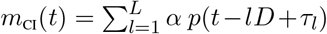. Here, *p(t)* is a rectangular pulse (pulse function) of unit amplitude and duration of 5 ms. Each rectangular burst was scaled by a factor *α* large enough to depolarize, on average, 50% of the MNs, triggering an action potential within the burst duration and up to 3 ms after its end. The 1/*D* rate (fundamental frequency) at which the depolarizing bursts occurred was varied between 0 and 4 Hz in 1 Hz steps across different simulations. To avoid that the bursts series were perfectly periodic, a random burst occurrence variability factor *τ*_*l*_ bounded between 0% and 20% of the corresponding mean inter-burst period was added. From an spectral point of view, this variability range implies that the harmonics of the burst frequency in the PSD have a smaller power than the fundamental burst frequency Lyons (2010). Each MN also received an *m*_II_(*t*), and a constant value *𝒞*_*j*_, as defined previously.

In additional non published analysis, we simulated a similar scenario but using a train of excitatory Dirac delta pulses as the iCI (instead of a train of rectangular bursts). The results were similar to those presented in this study.

#### cCI scenario

As mentioned in the introduction, the classic approach when simulating the firing properties of MN pools has been to consider cCI without impulsive CI. To assess this scenario, we ran simulations considering different inputs: (1) a sinusoid at a frequency matching the average DR of the simulated MNs (10.4 Hz, as in previous simulations, referred as ℱ_DR_); (2) a sinusoid at 20 Hz (simulating beta oscillations Ibáñez et al. (2021)); and (3) a sinusoid at 1 Hz.

To define these sinusoidal inputs, we generated independent realizations of white gaussian noise processes with zero mean and standard deviation of 1. We then band-pass filtered these realizations with 2nd order Butterworth filters centered at the three frequencies simulated and with 1 Hz bandwidth. We normalized this filtered noise by dividing it by its root mean square value. We then multiplied this signal by the squared root of the target power to achieve, resulting in *m*_CI_(*t*) = *α* cos(2*π*ℱ_i_*t*), with ℱ_i_ *∈* {ℱ_DR_, 20, 1} Hz. To determine the target power of this CI component, we used the maximum power that did not alter the coefficient of variation of the simulated force (*<* 0.5% of change) for the case of a cCI at ℱ_DR_. This target power was estimated in previous simulations. Each MN also received an *m*_II_(*t*), including a MN-specific constant value *C*_*j*_, as defined previously.

In an additional analysis of this scenario (second section in the results), we used a sinusoid at ℱ_DR_ with a power scaled to match that of the harmonic term at ℱ_DR_ of the iCI at 1 Hz. In this complementary analysis, the iCI did not exhibit variability in the bursts occurrence times.

#### Control scenario with no extra CI

To explore the baseline behavior of the MN pool in the absence of additional modulatory CI, we simulated a scenario in which the MNs did not receive any CI component (neither iCI nor cCI). This implies that the MNs only received *m*_II_(*t*) including the MN-specific constant, *𝒞*_*j*_, that made them fire at a mean dominant rate, ℱ_DR_, of 10.4 spikes/s with the variance previously reported.

#### Combined CI scenario

Here we simulated a scenario combining iCI and cCI. The objective was to analyze how impulsive inputs affect the ability of the MN pool to transmit cCI described in the current framework. We ran a set of simulations in which the MN pool received a CI formed by two different components: a sinusoid at either 20 Hz or 1 Hz, an iCI with a fundamental frequency of 1*/D* = 1 Hz. We used a cCI at 20 Hz because there exists evidence of the transmission of such kind of activity across the spinal MNs Bräcklein et al. (2022) Zicher et al. (2023). We used a cCI at 1 Hz to study the effects of the iCI in the frequencies related to the force control. The resulting common input becomes

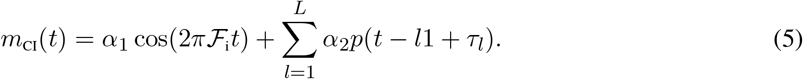

where ℱ_i_ is either 20 Hz, simulating the beta oscillations, or 1 Hz, simulating the force control signal. We tested different power levels corresponding to *α*_1_ to examine how the nonlinear effects triggered by the iCI depend on the strength of the cCI projection to the MN pool. Specifically, we varied the power in 10% increments relative to the maximum power used. This maximum was defined as the highest power level that did not alter the coefficient of variation of the simulated force by more than 0.5%, when using a cCI of 20 Hz. The iCI was defined as in the previous section. The power of each pulse, controlled by *α*_2_, was sufficient to depolarize a mean proportion of 50% of the MNs, as simulated previously. In our tests we compared results with and without including the iCI. In this context, we studied how the impulsive events affect the transmission of continuous signals at the known beta band and whether these effects depend on the power of the cCI component.

### Signal Analysis

The signal analysis include different blocks from preprocessing to the estimation of different metrics. Each block is described below.

#### Preprocessing of experimental dataset

First, for each subject, a data segment of 30 s corresponding to a period with a continuous constant 10% MVC force was extracted (this ensured that the decomposed MNs within that 30 s segment were firing steadily). The details about EMG decomposition are reported in the original article Ibáñez et al. (2021). Briefly, in the offline analysis, the HD-EMG signals were first band-pass filtered (20–500 Hz) using a second-order zero-lag Butterworth filter. The signals were then decomposed into motor unit spike trains MN_*j*_, using blind source separation techniques Holobar et al. (2014). Manual editing of the decomposed activity was conducted in accordance with previously described procedures Hug et al. (2021). The *i*-th spike time *t*_*i,j*_ of each *j*-th MN was defined as the onset of the motor unit action potential. To estimate this onset, double differential signals were computed from the monopolar HD-EMG recordings along the columns of the electrode grid, approximately aligned with the longitudinal direction of the muscle fibers.

For each MN decomposed we computed its instantaneous firing rate in non overlapping 1-s windows. MNs with a standard deviation in the instantaneous discharge rate larger than 1.5 spikes/s were discarded for subsequent analyses.

#### Power Spectral Density

For both simulated and experimental data, the PSD of the cumulative spike train (that is, the summed activity of all MNs) was computed using Welch’s method (MATLAB function *pwelch*) using 1 second windows with an overlap of 0.5 seconds and a frequency resolution of 0.25 Hz. The cumulative spike train was detrended before computing the PSD to remove the spectral component at 0 Hz. The simulated data included larger number of MNs than those obtained when decomposing experimental data. To compare experimental and simulated results, the cumulative spike train obtained from the simulated data was computed using 30 randomly selected MNs from the simulated data and this procedure was repeated 80 times, each with a different set of 30 simulated MNs, and the resulting PSDs were averaged. When computing the cumulative spike train in each iteration, we also computed the input-output gain at ℱ_DR_ averaged across iterations. Gain was defined as the ratio between the power at ℱ_DR_ of the cumulative spike train an the power at the same frequency in the common input *m*_CI_(*t*).

#### Intra Muscular Coherence (IMC)

The IMC is a widely used metric to estimate shared inputs across MNs within a pool Dideriksen et al. (2018). Since IMC is calculated by averaging the spectral activity of subpools of MNs across different time segments, it serves as a useful tool for estimating stationary inputs to the MN pool. In other words, the IMC is sensitive to inputs that influence MN activity homogeneously over time, such as cCI.

Briefly, to obtain the IMC, spectral coherence is computed between the activities of two equally sized MN subpools and the results are averaged across multiple iterations with different subpools of MNs. In this study, magnitude-squared coherence was computed using Welch’s method (*mscohere* MATLAB function), with 1 s windows, 0.5 s overlap and a resolution of 0.25 Hz. For the experimental data, we first defined two subpools of MNs with half of the total number of MNs available for each subject (this is the maximal number of MNs per subpool to compute the IMC). We built the cumulative spike train of each subpool, detrended them and computed the coherence. This process was performed in 80 iterations and the coherences for each iteration were averaged. In each iteration, the MNs belonging to each subpool were randomly selected from the total set of MNs. For the simulated data, 15 MNs in each subpool were randomly selected from the full simulated MN pool (177 MNs) in each iteration (with the two permuted subpools, this implies a total pool of 30 MNs at each iteration). The same procedure was then applied to compute coherence across 80 iterations and the results were averaged.

#### SPIKE distance

The SPIKE distance Kreuz et al. (2013) is a metric to quantify synchrony between spike trains for both simulated and real data.The SPIKE distance is a time-resolved measure of dissimilarity between spike trains that focuses on the precise timing of spikes rather than overall spike counts or firing rates. At each moment in time, it assesses how close the nearest spikes are across spike trains by considering both the preceding and following spikes. These instantaneous differences are then normalized by the local inter-spike intervals, allowing the metric to become independent of variations in firing rates and providing a sensitive, temporally detailed comparison of spike timing. Details can be found in Kreuz et al. (2013), but the formula for the time-varying SPIKE distance between two spike trains (here, *c*_*n*_(*t*); *n ∈* {1, 2}) is computed by defining the timing, 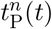 of the preceding spike to time instant *t* in the spike train *n* as

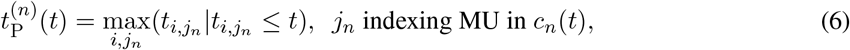

and the time instant following time t as

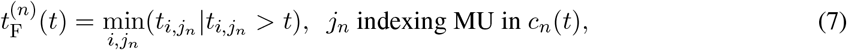

from where an interspike interval from later preceding to earliest following spikes at time *t* can be defined as

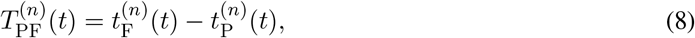

and the difference between spikes timings at different sub pools for the preceding and following ones can be defined as

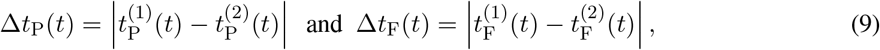

respectively. This definitions lead to propose a dissimilarity spike trains as the average of the inter pools spike differences normalized by the mean interspike distance.

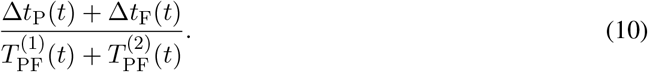

Aiming to be more local in time, giving more weight to inter pools spike differences closer to analyzing time *t*, defining

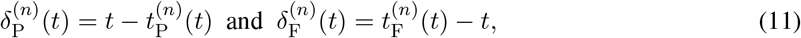

and weighting inversely to the across subpools mean of these distances, the following dissimilarity measure is obtained Kreuz et al. (2013).

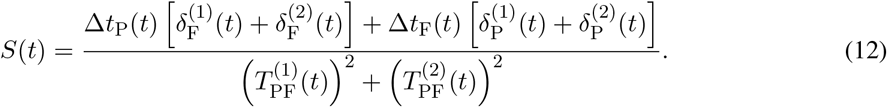

The SPIKE distance *S*(*t*) is a time-varying metric that can be computed in a multivariate way, by averaging the spike distances of each pair of spike trains. The SPIKE distance is a dissimilarity index: when the MNs are more synchronized, the index shows smaller values, bounded to the interval [0, 1]. To implement the SPIKE distance in MATLAB, we used the code developed in Kreuz et al. (2013).

We wanted to relate the SPIKE distance to the population activity of the MN pool. As stated in the introduction, individual MNs are governed by their DR. A synchronized pool of MN is also partially dominated by the DR of the pool, as the MNs are firing simultaneously. This implies that spectral power at ℱ_DR_ serves as an indicator of the pool’s synchrony. Therefore, we calculated the SPIKE distance at time points corresponding to increases in the DR-band power of the cumulative spike train. This measure of synchronization based on the power at the ℱ_DR_ was applied to all three simulated scenarios and the experimental data, allowing for a comparison to identify which scenario most closely resembles the recorded activity.

To analyze this, the cumulative spike train was band-pass filtered (2nd-order Butterworth, 4 Hz bandwidth centered at ℱ_DR_), and the instantaneous amplitude was extracted using the Hilbert transform. Time points where the amplitude exceeded the mean plus one standard deviation were identified as synchronization events. Centered around each *k*-th event timing *t*_*k*_, the time of maximal instantaneous amplitude at the filtered cumulative spike train, a 2 s window was extracted, and the SPIKE distance was averaged within that *k*-th window as an estimate of MN synchronization during periods of elevated power of the cumulative spike train at ℱ_DR_.

As before, for the simulated data, 30 MNs were randomly selected in each iteration, and the SPIKE distance was computed for that subpool. This process was repeated over 80 iterations. In each iteration, the SPIKE distance was averaged across all the *k*-th windows centered on synchronization events and across the random time windows. These values were then averaged across iterations. For the experimental data, the same procedure was applied using all decomposed MNs for each subject.

To express the magnitude of the synchronization event, we computed the minimal value of the averaged SPIKE distance. We picked the minimal SPIKE distance value because the SPIKE distance is a normalized measure (bounded to the interval [0, 1]) and thus can be compared across conditions. To assess whether the experimentally observed SPIKE distance truly resulted from the influence of the described iCI rather than from the intrinsic statistical properties of the data, we conducted an additional analysis using surrogate data. For each subject, we generated a surrogate dataset by circularly shifting each spike train by a random duration between 2 and 8 seconds, drawn from a uniform distribution. SPIKE distance was then computed following the same procedure as with the original data (averaging values within the windows centered around peaks in the filtered cumulative spike train). This process was repeated over 100 surrogate datasets per subject, with the resulting SPIKE distances averaged across iterations. If the reduced SPIKE distance (increased synchrony) in the experimental data were solely due to the intrinsic statistical properties of the dataset, then the minimum SPIKE distance observed in the surrogate data should closely match the experimental value. Conversely, if the reduction reflects the influence of an underlying input, a clear difference between the minimum surrogate and experimental values should be expected.

#### Pearson Correlation coefficient

The correlation coefficient was used to quantify the cCI transmission across the MN pool in the final part of the study (where simulations combining impulsive and sinusoidal inputs were used). To account for potential phase shifts or lags between the signals, we computed the cross correlation function between the CI and the filtered cumulative spike train and select the maximal value of this function at each iteration. In each simulated scenario, the correlation was computed using cumulative spike trains composed of either 10, 30 or 177 MNs, to assess transmission when considering smaller subpopulations versus the entire MN pool. For the cases of 10 and 30 MNs, we performed a total of 80 iterations. At each iteration, the MNs were randomly picked from the entire pool and the spike trains of the selected MNs were summed. The resulting cumulative spike train was band-pass filtered in the same frequency band as the frequency used to generate the cCI using a 2nd order Butterworth filter of 1 Hz bandwidth centered at the frequency of interest. The correlation coefficients (maximal value of the cross correlation function) were averaged across iterations.

#### Statistics

The central premise of this paper is that MNs do not receive information solely in the form of cCI. To determine whether the experimentally observed magnitude of the synchronization events is significantly greater than that observed under the influence of cCI at different frequencies, or in the absence of any inputs (reflecting the intrinsic behavior of the simulated pool), two-sample t-tests were performed between the minimal values in SPIKE distance for each subject and that of each simulation condition. The Normality of the data was assessed visually by qqplots and tested statistically by Lilliefors tests. In the cases where the data was not normally distributed (as determined by the Lilliefors test), Wilcoxon rank sum tests were used (in comparisons between the experimental data and both the cCI at ℱ_DR_ and the no-CI condition). A significance level of 0.05 was used for t-tests and rank sum tests, with all p-values adjusted for multiple comparisons using the Bonferroni correction. All the results are reported as mean ± standard deviation.

## Results

This section is divided in three parts. First we compare the experimental data with simulated scenarios to look for evidence supporting the hypothesis that MN pools receive iCI. Then, in the second and third parts, we show how iCI affect the transmission of cCI by MNs and how this may limit the estimation of these inputs from MN outputs. For the comparison of the metrics with the experimental data and for the transmission analysis (Sections 1 and 3 of the results), we used the case of iCI having a dominant rate 1*/D* = 1 Hz (1 burst per second), as this was approximately the mean rate observed in the experimental data.

Regarding the experimental data, the average number of MNs per subject included in the analysis was 23.9 ± 6.4. The average firing rate ℱ_DR_ of the MNs per subject was 10.3 ± 1.2 spikes/s.

### Simulations considering Impulsive Common Inputs reproduce key characteristics of MN pool activity recorded experimentally

An example of the simulated MN activity in the scenario using depolarizing iCI is shown in Fig. 2 A top. The figure shows how MNs synchronize their firings every time an impulse occurs. As a consequence of this synchronization, the amplitude of the summed MN activity filtered at ℱ_DR_ increases briefly (Fig. 2 A bottom). This increase in power at ℱ_DR_ reflects that the MN pool is momentarily behaving more similarly to a single MN (where the power spectrum is dominated by a peak at the fundamental frequency, that is, ℱ_DR_). Similar dynamics may be qualitatively observed in experimental data, with sporadic alignments of MN firings associated with increases in the power of the cumulative spike train in frequencies around ℱ_DR_.

**Figure 2.**
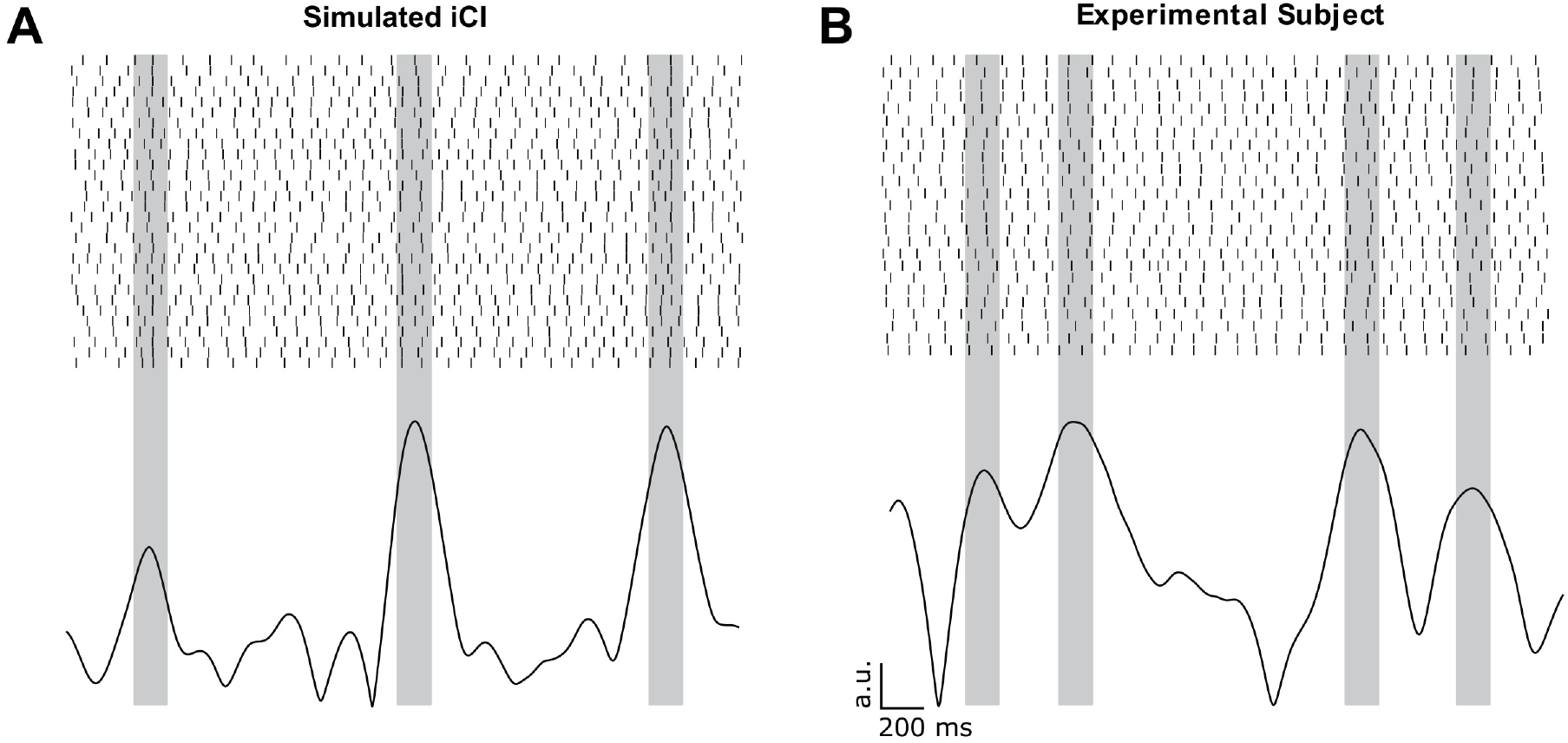
**A:** Raster Plot (3 seconds) and instantaneous amplitude (obtained with the Hilbert transform) of the cumulative spike train of a simulated population of 30 MNs receiving an iCI of 1 burst per second, synchronizing its firings at the times of the bursts. **B:** Raster Plot (3 seconds) and instantaneous amplitude (obtained with the Hilbert transform) of the cumulative spike train of a s population of 25 MNs from a representative subject. The plots show how the peaks of the filtered cumulative spike train indicate an alignment of the MN firings.

Based on the above reasoning, in order to identify moments of synchronization (reflecting iCI), we looked for the time instants at which the power at the average ℱ_DR_ of the summed MN activity was high (more than one standard deviation above the mean) and located the peak times of these events. For the experimental data, the average rate of the detected events with high power at the ℱ_DR_ frequency was 0.71 ± 0.08 events/s. When using the surrogate datasets, we did not observe a substantial difference in the number of detected events (0.69*±*0.02) with respect to the true dataset. However, the magnitude of the surrogate SPIKE distance was much less pronounced relative to that of the true dataset (discussed below). In the case of simulations using iCI, the mean rate was 0.87 ± 0.11 events/s. In the cases where simulations did not include cCI or they did include cCI, the rates were 0.64 ± 0.11 events/s for the cCI at ℱ_DR_; 0.78 ± 0.04 events/s for the cCI at 20 Hz; 0.73 ± 0.13 events/s for the cCI at 1 Hz; and 0.73 ± 0.01 events/s for the no-CI scenario.

Fig. 3 shows the MN synchronization profiles (measured with the SPIKE distance metric) around the events of high power at ℱ_DR_. The first panel in the figure shows the results obtained from experimental data (Fig. 3 A), both for the true recorded data (averaged across subjects, blue curve, with grey curves representing individual subjects) and for the surrogate data (averaged across subjects, yellow curve). For the true experimental data (blue curve), the synchronization profile presented a short-lasting marked reduction (negative peak) around 0 s, reflecting that the moments of high power at ℱ_DR_ are associated with a sharp and brief increase in spike synchronization across MNs, as predicted. When performing the surrogate approach (by circular shifting each spike train by a factor between 2 and 8 seconds and averaging across 100 iterations per subject), the average minimal SPIKE distance obtained was 0.286, significantly higher than the average minimal SPIKE distance of the true data, 0.255 (*p* = 1.12 *×* 10^*−*7^). These results indicate that the synchronization observed in the experimental data (blue curve) cannot be attributed to the intrinsic statistical structure of the dataset.

**Figure 3.**
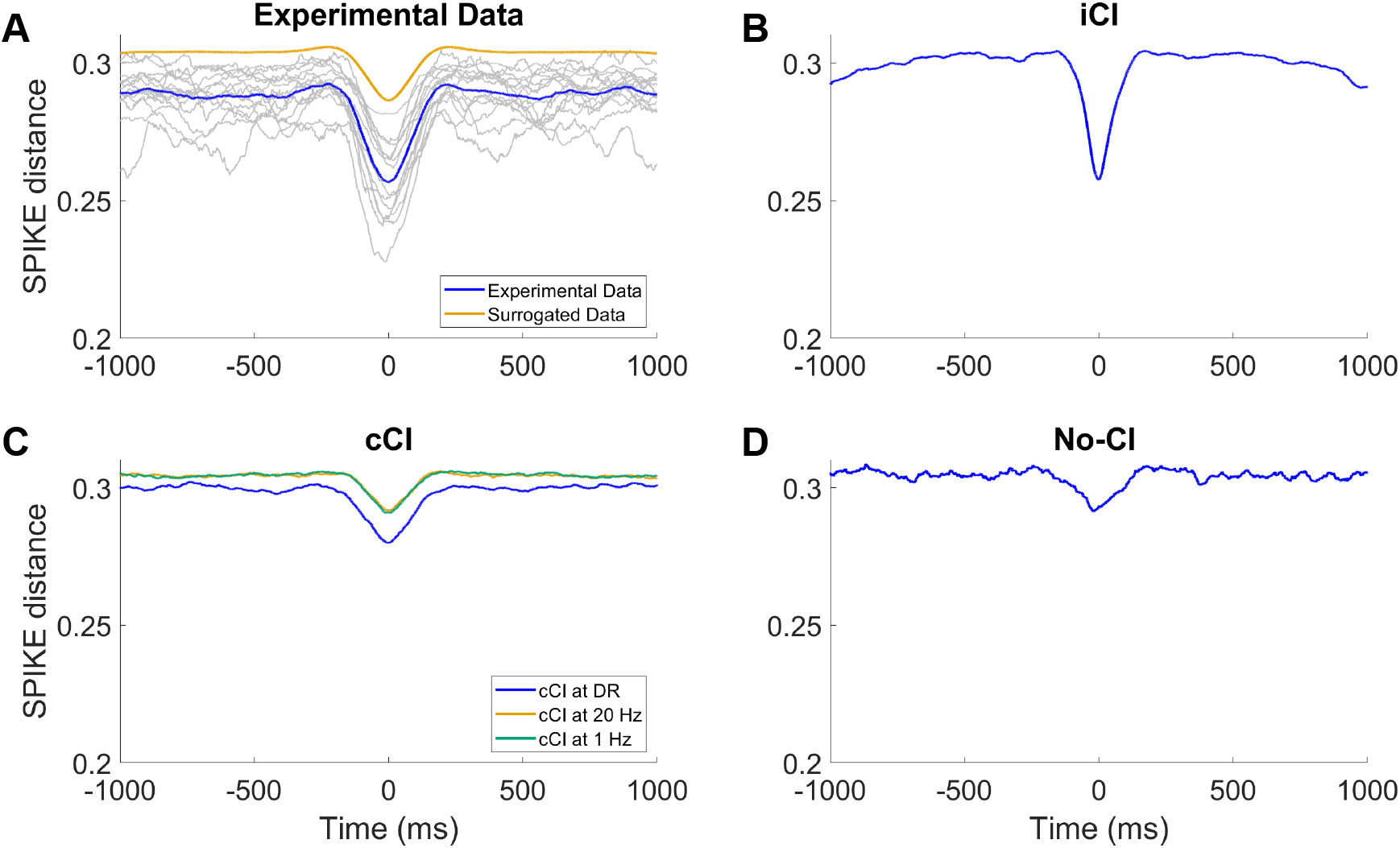
Average SPIKE distance changes in segments around time points where the power of the summed MN activity presents a maximum at ℱ_DR_. The panels show the results obtained using experimental data (grey curves representing individual subjects and colored curves representing mean of the true data, blue, and of the surrogate data, yellow), and simulations with iCI, cCI and no-CI. The figure shows a clear resemblance in MN synchronization between experimental data and simulations using iCI. The other scenarios do not lead to comparable results, as they do not show a clear evidence of synchronization changes linked with increases in power of the summed MN activity at ℱ_DR_.

Fig. 3 B represents the SPIKE distance from simulated data using iCI (rate of 1 burst per second), averaged across simulations. Interestingly, this scenario also reproduces the short-lasting marked reduction (negative peak) around 0 s and overall shows a very similar pattern than that observed in the experimental data (blue curve in panel A). In particular, the SPIKE distance for the simulated iCI showed an average minimal value of 0.256 not significantly different from the minimal values obtained in the experimental case (*p* = 0.85). This implies that iCI simulations induced a synchronization across MNs that closely matched the experimental observations.

Simulations only considering cCI (Fig 3 C, curves averaged across simulations) did not lead to synchronization events at around 0 s that were comparable to the previous two cases. When cCI were simulated, the average minimum SPIKE distance level was always significantly greater than for experimental data, regardless the frequency of the input. Specifically, the average minimum SPIKE distance reached in each case was 0.278 for the cCI at ℱ_DR_ (*p* = 3.96 × 10^−4^); 0.291 for the cCI at 20 Hz (*p* = 5.13 *×* 10^*−*7^); and 0.290 for the cCI at 1 Hz, (*p* = 7.80 × 10^−7^). Similarly, when simulating the no-CI scenario (Fig 3 D, curve averaged across simulations), the average minimal SPIKE distance was 0.291 (also significantly greater than the value experimentally obtained, *p* = 2.41 × 10^−5^).

Overall, these results indicate that cCI or the mere variability of MN activity in the absence of CI cannot account for the observed sporadic and brief synchronization events observed experimentally. iCI, on the contrary, can reproduce the experimental results successfully, supporting their plausibility as an underlying input structure.

To further characterize the effects of iCI on MN activity at the population level, we computed and compared the PSD and IMC functions obtained from experimental data and the simulations. The PSD and IMC functions are typically used to analyze the population activity of MNs as they measure the spectral distribution of the signals that are sampled by the pool. Fig. 4 A-B shows the average PSD and IMC obtained from the experimental data. These plots are qualitatively similar to those reported in other studies in the literature Farina and Holobar (2016); Ward et al. (2013) and show two characteristic features. The PSD is dominated by a peak at the frequency matching the average ℱ_DR_ of the MNs (this is typically around 10 spikes/s in the tibialis anterior muscle for low-force contractions as in this study). This peak is however not present in the IMC, indicating that the MN pool does not contain an oscillatory common signal at ℱ_DR_ that dominates the spectra of the MNs. The same analysis using the simulated data with iCI with a rate of 1 Hz (Fig. 4 C-D) leads to results that are very similar to those observed experimentally, with a relatively low IMC level at frequencies around ℱ_DR_ despite the presence of a prominent peak at that same frequency in the PSD. This is likely related to the induced transient synchronization events of MNs at the times of the impulses. Due to the short-lived nature of these events, the IMC is not affected by them and shows a lack of common contents in the MNs at ℱ_DR_ (Fig 4 D).

**Figure 4.**
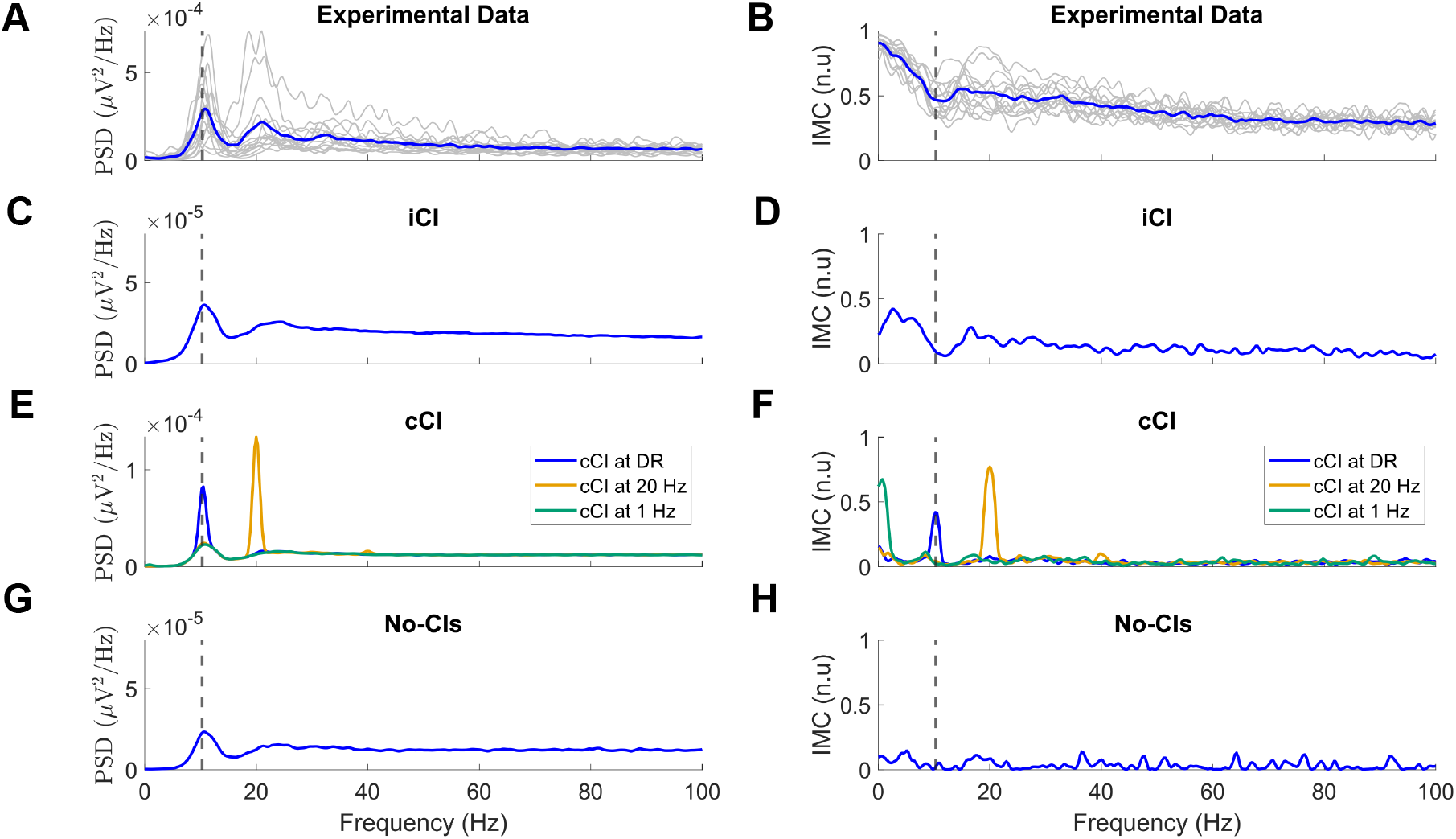
PSDs and IMCs of the summed activity of MNs for experimental data (A-B), simulations using iCI (C-D), simulations using cCI (E-F) and simulations that do not include any CI (G-H). Curves are averaged across subjects or simulations. The vertical dashed line indicates the mean discharge rate of the pool.

Simulations using cCI (Fig 4 E and F) lead to PSD and IMC functions that were similar in shape to the power spectrum of the CI itself (as expected due to the capacity of MN pools to linearly amplify cCI) Farina et al. (2014); Farina and Negro (2015). In these cases, a prominent peak at the frequency of the CI can be observed in the IMC, which implies that in the presence of only cCI, the combined activity of MNs should tend to be dominated by contents related to the inputs. Finally, the scenario in which no CI are simulated leads to values of the IMC close to 0 at all frequencies, which is expected given the absence of common signals across the MNs (Fig 4 H).

In summary, we observe relevant similarities between the experimental data and simulations using iCI. These similarities are not reproduced when simulations do not include impulsive inputs but instead use cCI. These results strongly suggest that the MN pool receives at least part of its drive in the form of intermittent iCI. The next sections address how these impulsive inputs can affect the linearity assumptions typically made in the interpretation of signals sampled by MN pools.

### Impulsive inputs alter the linearity of MN pools

Under the influence of cCI, MN pools act as linear amplifiers of these inputs Farina et al. (2014). However, in the case where iCI also drive individual MN activity, the repeated transient synchronization across MNs (once every time an impulse is produced) may cause the linear behavior of the system to be altered. To assess this, we conducted several tests using the simulations involving iCI.

First, we assessed how the fundamental frequency of the iCI (that is, the average rate at which impulsive events occur) affects the power spectrum of the summed activity of MNs. In line with the previous section, 30 MNs were used to obtain the PSD of their summed activity (randomly picking 30 different MNs several times and averaging the obtained results). If the iCI do not affect the linear behavior of the MNs, it should be expected that the PSD of the summed MN activity is only modified according to the spectral contents of the impulsive input (a linear transmission). However, we observe that this is not the case when iCI are simulated. Fig. 5 shows the evolution of the PSD of the summed MN activity when the rate of occurrence of the impulsive inputs increases from 0 events/s to 4 events/s. As the rate of impulsive events increases, the PSD average level increases although, relevantly, the PSD level at ℱ_DR_ increases at a higher speed than the rest of the frequencies. In fact, we observe that the power at ℱ_DR_ in the cumulative spike train increases monotonically when the rate of impulsive events increases despite the gain at that frequency shows a very different trend (inset in Fig. 5 shows that the gain at ℱ_DR_ does not follow a linear increase with the input frequency). Such an outcome is inconsistent with the predictions of a purely linear system, where the output power at ℱ_DR_ would be expected to scale proportionally with the input power at the same frequency.

**Figure 5.**
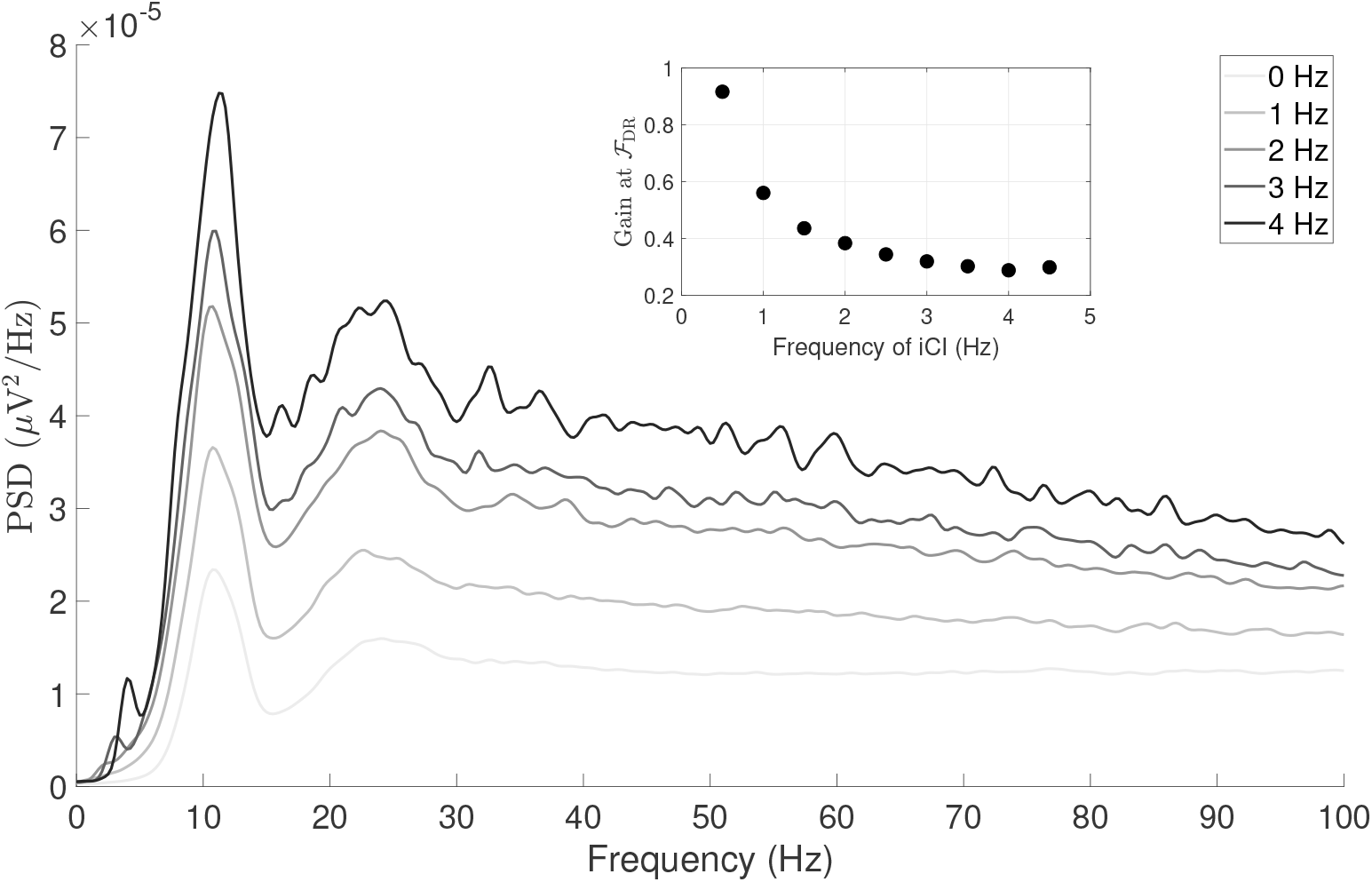
Spectrum of a pool of 30 MN receiving iCI from 0 Hz (no bursts) to 4 Hz, in 1 Hz increments. The inset figure represents the gain at ℱ_DR_ with respect to the bursts frequency.

Since the linear behavior of the MN pool may depend on the number of MNs considered, we conducted a second test where we compared the gain (output power divided by the input power) at ℱ_DR_ for different numbers of MNs when the CI was either an iCI producing one impulsive event per second or a cCI modeled as a pure sinusoid with a frequency ℱ_DR_ and with a power at that frequency matched to the power of the iCI (that is, both CI had the same power at ℱ_DR_ but different power at other frequencies) (Fig. 6 A). The gain at ℱ_DR_ of the summed activity of MNs increased with the number of MNs considered in both scenarios (Fig. 6 B). However, while the power at ℱ_DR_ of the summed MN activity increased linearly in the case of the cCI, the trend became exponential with the iCI (*R*^2^ = 0.97 of fit for an exponential model of the form *y* = *a* · *e*^*b·x*^, with parameters *a* = 0.60 and *b* = 0.01, being *y* the gain at ℱ_DR_ and *x* the number of MNs included) and the output power at ℱ_DR_ diverged from the power obtained with the cCI as more MNs were considered. This reflects the deviation from the linear behavior when the impulsive input is used (two input signals with the same power at a certain frequency lead to output signals with different power at that same frequency) and it also reflects that the non-linear effects produced by the MN pool do not decay when larger pools of MNs are considered. The spectra of the cumulative spike train for both input scenarios across the different number of MNs evaluated are shown in Fig 6 C (cCI) and D (iCI) respectively. In both spectrums, it can be observed that the peak at ℱ_DR_ increases more rapidly with the number of MNs in the iCI case (Fig 6 D).

**Figure 6.**
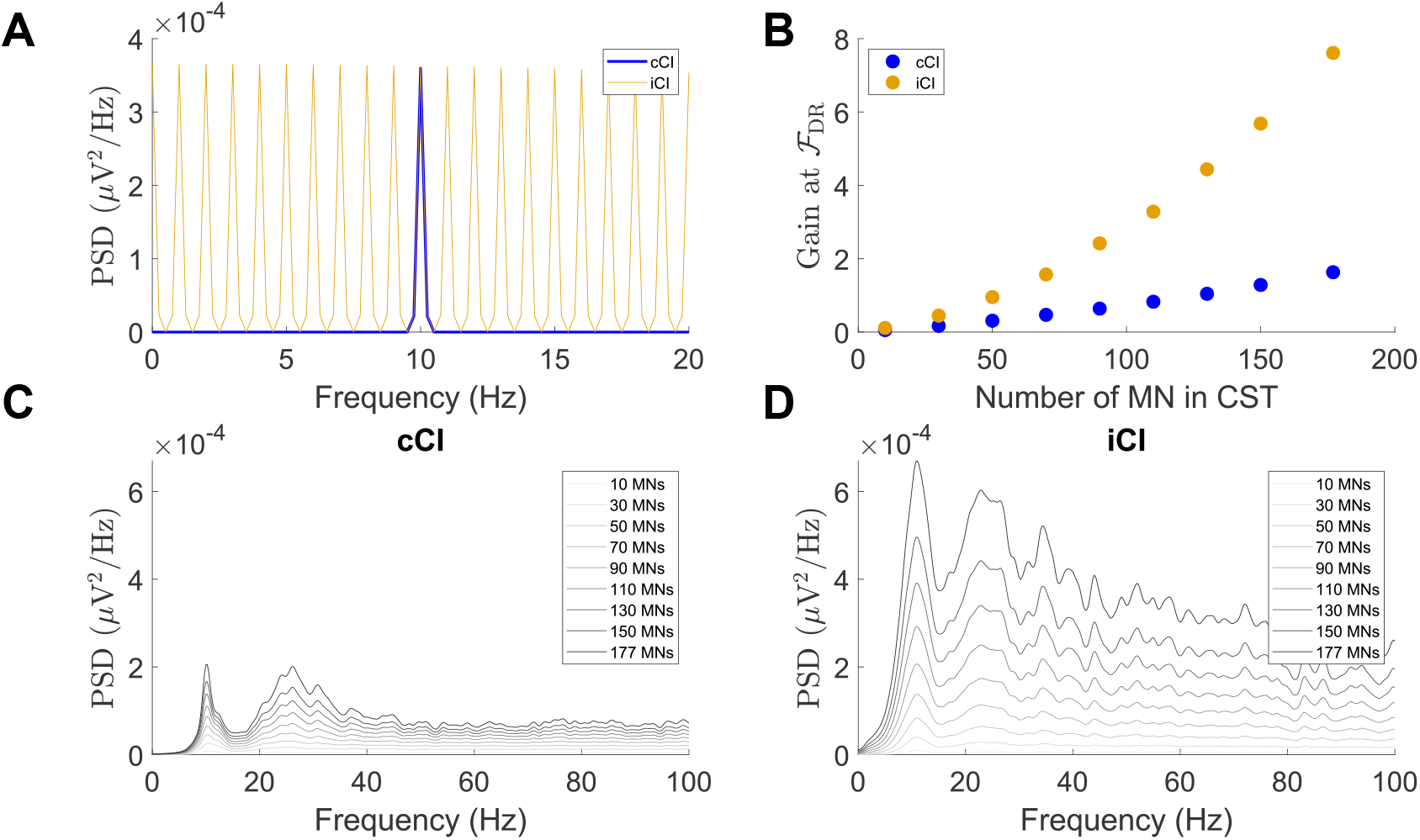
**A:** Spectrum of two types of CI simulated as a pure tone (blue) or a burst train (yellow) with 1 Hz fundamental frequency **B:** Ratio (gain) between the output and input power at the average ℱ _DR_ of the simulated MNs when the CI is a pure tone (blue) or a burst train (yellow). Each point correspond to the average across 10 simulations **C:** PSD of a MN pool receiving a sinusoid at ℱ_DR_ Hz, evaluated for different number of MNs in the cumulative spike train **D:** PSD of a MN pool receiving a iCI at 1 Hz frequency, evaluated for different number of MNs in the cumulative spike train

In summary, this subsection shows that the presence of iCI can alter the behavior of MN pools as linear amplifiers of CI. The next subsection analyzes how this effect can deteriorate the estimation of cCI in the presence of iCI driving MN activity.

### Impulsive input-induced synchronization alters signal transmission by the MN pool

In this section, we examine how iCI can affect the transmission of cCI by the MN pool. The main objective in this analysis is to assess how reliably cCI in the beta band or in the low-frequency band (related to the force control) can be decoded from MN activity when the MN pool is also driven by iCI, which disrupt its linear transmission properties as previously described.

To assess this, in different sets of simulations we evaluated the transmission of cCI at 1 Hz and 20 Hz by a MN pool in two different conditions: the MN pool receiving only cCI or receiving both cCI and iCI. We tested increasing power levels for this cCI (increasing levels of signal to noise ratio), with the maximum corresponding to the maximal power that did not significantly influence the coefficient of variation in the simulated force, as detailed in the Methods. For each power level, we tested how it was transmitted by the MN pool with and without an additional iCI, evaluating this for 10, 30 and 177 MNs in the cumulative spike train generated to estimate the input from the output of the MNs. The impulsive input was generated with a fundamental frequency of 1 Hz and the same power as in previous sections. To assess the transmission of the cCI, we measured the maximal value of the cross correlation function between that input and the cumulative spike train of the MN pool filtered around 20 Hz or 1 Hz (depending on the frequency of the simulated cCI). For this analysis, the MN pool was firing at an average rate of approximately 10.4 Hz, in line with previous sections.

Fig. 7 summarizes the results obtained in these simulations. The main finding is that the presence of iCI consistently reduces the correlation between the cumulative spike train and the cCI, regardless the number of MNs included, the frequency of the cCI or the input power level. However, this reduction in the correlation was more pronounced for lower powers of the CI (low signal to noise ratios), as expected.

**Figure 7.**
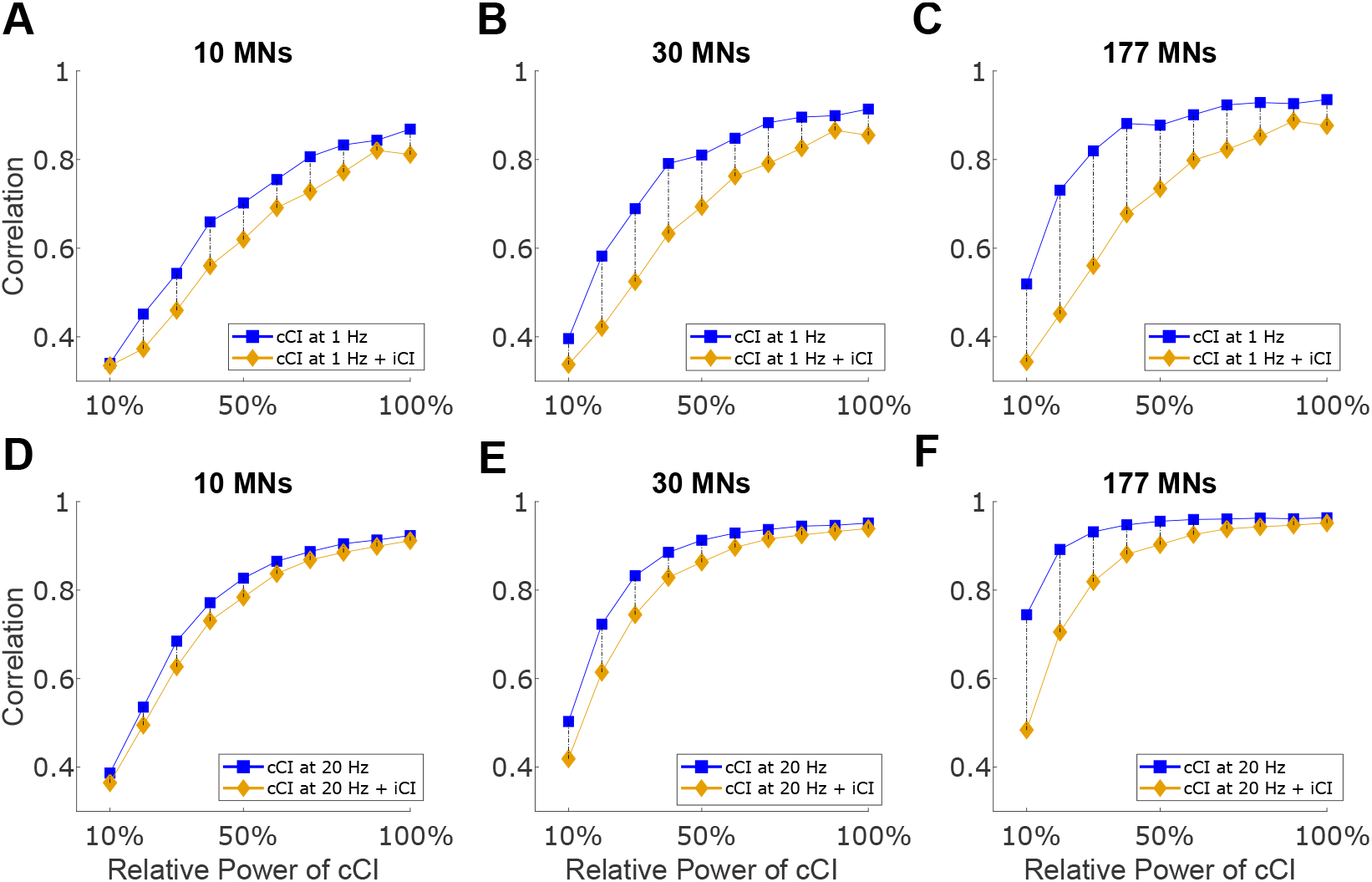
Correlation between the filtered cumulative spike train and a cCI at 1 Hz (A,B,C) or 20 Hz (D,E,F). The cumulative spike train was obtained summing the activity of 10 MNs (A and D), 30 MNs (B and E) or 177 MNs (C and F) and then band-pass filtering the result around 1 Hz or 20 Hz (depending on the simulated input). The black dashed lines highlight the correlation difference for each power level used for the cCI. The power of the cCI is expressed as a percentage of the maximal power used (maximal power such that it does not modify the coefficient of variation of the simulated force).

Two main findings can be identified from these simulations. First, the correlation drop caused by the presence of iCI becomes more pronounced with increasing numbers of MNs (the difference between the blue and the yellow curves is more pronounced when more MNs are included). This is in line with Fig 6 B, which shows that increasing the number of synchronized MNs in the cumulative spike train amplifies the non-linear behavior of the pool. Second, the correlation drop observed at 1 Hz (Fig 7 top row) is greater than that at 20 Hz (Fig 7 bottom row) for all MN group sizes evaluated, indicating that the disruptive effects of the iCI are more pronounced at lower frequencies.

It is important to remark that the impulsive events in these simulations appeared only once per second, while the correlation was computed over 30-second simulations. This indicates that the distortion caused by each pulse was brief and localized, allowing the MN pool to recover linear transmission properties during the remaining time, resulting in the high overall correlation values obtained in the simulations.

Overall, these analyses indicate that iCI impair the MN pool ability to sample and linearly transmit cCI. This outcome is consistent with the fact that impulsive inputs push the MN pool toward a nonlinear behavior. The extent of this disruption depends on the balance between the power of the cCI and the power (or frequency) of the iCI: the weaker the cCI, the more pronounced the effect of the impulses on its transmission, and conversely, stronger cCI are more resilient to the disruptive effects of the iCI.

## Discussion

Studying the population activity of spinal MNs is essential for understanding the neural strategies underlying muscle control and force generation. Many motor neuron models account only for continuous inputs characterized by relatively slow dynamics Farina et al. (2014); Farina and Holobar (2016); Zicher et al. (2023); Watanabe and Kohn (2017). However, within this framework, certain features of experimentally observed MN activity cannot be accurately reproduced. In this study, we propose and analyze the existence of common impulsive signals transmitted to the MNs that force a temporal synchronization of their action potentials. Using both simulations and experimental data from healthy human subjects, we indirectly showed evidence for the first time that MN pools can receive significant influences through impulsive common inputs (iCI) that alter their linear behavior. In this context, we use the term impulsive implying that these neural signals that drive MN activity occur during periods of time that are much shorter than the spiking frequency of the driven MNs. The presence of such iCI can successfully replicate certain aspects of the MN activity not explained by simulations that only include continuous CI (cCI), such the common spectral contents across MNs and the spiking synchronization at the population level. These findings have important relevance in the current understanding of motor control and have an impact on the development of neural interfaces that extract information from the activity of populations of MNs. In the following sections, we discuss the potential role of these impulsive inputs and their possible origins.

### Potential role of the impulsive inputs on the motor pathway

Motor control models commonly assume that the motor cortex exerts continuous control over muscles Shadmehr and Wise (2004). However, it has also been proposed that higher-level motor commands may be intermittent Karniel (2013). One compelling idea is that the motor system decomposes complex or prolonged actions into discrete submovements that serve as building blocks for full motor behaviors. Such segmentation could arise from a limited temporal planning horizon, requiring ongoing updates during execution, as suggested by computational theories Guigon (2023). Although submovement-rate modulation is often associated with slow or prolonged movements, it may apply to a much wider range of motor behaviors.

Coordinating these submovements might require intermittent control signals—such as impulsive bursts—that synchronize a substantial proportion of the active motor neurons in a pool. These synchronized events could act as neural “reset” points, aligning the system for the initiation of the next submovement. Beyond preparation, such bursts might also serve a sensory–motor monitoring role, allowing the system to reassess the state of the muscles even during steady or constant tasks.

The presence of iCI could have measurable consequences for movement. Low-frequency components of the cumulative spike train strongly correlate with the force output of a muscle Thompson et al. (2018). Given the low rate of impulsive events observed here (typically less than one per second), their influence would be expected primarily in the low-frequency range of motor neuron activity and force output (1–2 Hz) Christou et al. (2004, 2003); De Luca et al. (1982). Indeed, it is well established that the CNS struggles to maintain perfectly stable force Enoka and Farina (2021), and force fluctuations at target levels above 10% of the maximal voluntary contraction appear largely driven by variability in the common modulation of motor unit discharge times Negro and Farina (2012). In this framework, iCI could contribute to force variability by injecting power into the 0–2 Hz band, thereby increasing signal variance and reducing steadiness compared to a system without such input.

It is also possible that impulsive inputs have no direct functional role. They may simply reflect incidental activity originating from other brain regions—such as the motor cortex—that reaches spinal motor neuron pools because the nervous system has not evolved mechanisms to suppress them. In such a scenario, the occurrence of iCI would be a byproduct of neural architecture rather than an adaptation. For example, synchronized bursts could arise from passive attentional processes triggered by salient external stimuli, such as a loud sound Novembre et al. (2018), leading to transient excitations in motor cortical areas that propagate downstream to spinal motor neurons without functional filtering.

### Possible origin of the impulsive inputs

Due to the nature of our experimental dataset and the limited access to spinal MNs, it is difficult to accurately characterize the properties of these impulsive inputs, such as their strength, periodicity, or innervation pattern, among other features. Furthermore, the absence of simultaneous recordings from other regions of the nervous system makes it challenging to speculate on the potential sources of these impulsive inputs. One possible source for this activity is the motor cortex, as there exist evidence in the literature suggesting the existence of short-lived events that can be sent to the MN pool, such as the beta bursts Echeverria-Altuna et al. (2022); Bräcklein et al. (2022) or drastic changes in cortex activity related to sequences of inhibition and excitation, modulated by external stimuli Novembre et al. (2018). The reticular formation is also a structure to be considered in this context. The reticulospinal tract has been described as one of the main sources of neural drive during voluntary control of movements Glover and Baker (2022). Interestingly, the reticulospinal tract has also been proven to mediate the acceleration of reaction time when a loud sound accompanies the cue Tapia et al. (2022). This suggests that the reticulospinal tract is able to generate additional inputs to the MN pool that modulate voluntary movement, in addition to the commands controlled by the motor cortex.

Future research analyzing the particular role of these superior structures during the tracking of different forces will provide further clarity about the generation of these intermittent events.

## Acknowledgements

JYM, PL & JIP were supported by the European Research Council (ERC) under the European Union’s Horizon Europe research and innovation program (ECHOES project; ID - 101077693).J.I. was supported by MICIU/AEI and FEDER, UE (Grant PID2022-138585OA-C32) and by a consoli dacio’n investigadora grant (CNS2022-135366) funded by MCIN/AEI/10.13039/ 501100011033 and UE’s nextGenerationeU/PRTR funds. A.P-V was supported by the European Union’s Horizon Europe research and innovation programme under the Marie Sk^3^odowska-Curie grant agreement No 101151398. Computa tions were performed using ICTS NANBIOSIS (HPC Unit at University of Zaragoza). DF was supported by the EPSRC under the project NISNEM Technology (EP/T020970/1) and by the ERC under the Synergy Grant project Natural BionicS (810346).

## Code Availability

The computational model and analysis code used in this study were implemented in MATLAB. The code is available upon request from the corresponding author.

